# Oral Microbiota Composition in Children and Adults During Spanish COVID-19 Lockdown: Impact of Home Self-Confinement and SARS-CoV-2 infection

**DOI:** 10.1101/2025.09.30.679491

**Authors:** Miguel Blanco-Fuertes, Gerard González-Comino, Pedro Brotons, Aleix Lluansi, Rosauro Varo, Desiree Henares, Cristian Launes, Maria Cisneros, Mariona F de Sevilla, Juan J. Garcia-Garcia, Alex Mira, Quique Bassat, Carmen Muñoz-Almagro

## Abstract

**Background:** The COVID-19 pandemic changed society’s habits and customs due to the social restrictions and health measures imposed during the first half of 2020. This study analyzes the composition of the oral microbiota in relation to age, household cohabitation, SARS-CoV-2 infection, and COVID-19 severity among children and adults under home confinement in Barcelona, Spain.

**Methods:** A prospective study conducted involving children and adults confined during the COVID-19 pandemic in the Barcelona Metropolitan Area between April and June 2020 included multiple cases of several participants living within the same family household. Saliva samples were collected from all participants, and microbiota composition was characterized through 16S rRNA gene sequencing.

**Results:** A total of 142 adults and 265 children living in 121 family households were included in the study. All 142 adults had a prior confirmed SARS-CoV-2 infection, and 20 (14.08%) of them had a history of severe COVID-19. SARS-CoV2 infection was detected in 58/265 (21.89%) of children; all of them were asymptomatic. Oral microbiota composition and diversity did not differ by SARS-CoV-2 infection status in children. In contrast, adults with severe COVID-19 exhibited lower microbiota diversity and distinct microbiota composition compared to those with mild disease symptoms. Age-related differences in oral microbiota composition were marked in the younger children groups. Additionally, cohabiting individuals shared more Amplicon Sequence Variants (ASVs) than non-cohabitants.

**Conclusions:** Age and cohabitation strongly influenced oral microbial composition. Our study demonstrates that oral microbiota composition in adults varies according to COVID-19 severity, whereas such microbial shifts are not observed in asymptomatic pediatric populations, regardless of infection status.

## Background

Severe acute respiratory syndrome coronavirus 2 (SARS-CoV-2), the etiological agent of the coronavirus disease 2019 (COVID-19), was first identified in patients with pneumonia of unknown origin in December 2019 in Wuhan, China (1,2). As a result of its global spread, the World Health Organization (WHO) declared the SARS-CoV-2 outbreak a pandemic in March 2020 (3). Since the first case of COVID-19 was reported in Spain, the number of new cases increased exponentially over time. In response, the Spanish government declared a state of emergency from March to June 2020 to contain the spread of the disease and prevent the health system from collapsing. Consequently, a series of measures were implemented, including strict home confinement, the closure of non-essential businesses, and the suspension of face-to-face educational, cultural, and recreational activities (4).

The field of human microbiome research has led to a more comprehensive understanding of the microorganisms that inhabit the human body and their role in health and disease (5,6). Among other insights, it has provided a better understanding of diseases caused by pathobionts, which are microorganisms that asymptomatically colonize the human body but can also cause severe diseases under certain conditions (7). It is known that the human microbiome exhibits high variability across different body sites and between individuals, reflecting the multitude of variables that can influence its composition (8). The COVID-19 pandemic and the resulting lifestyle changes have had a major impact on the health of the population. However, little is known about how it has affected the human microbiome.

The oral cavity is home to one of the most diverse and abundant microbial communities in the human body (5), whose equilibrium is responsible for the maintenance of a healthy state (9). One of the most commonly used samples for studying the oral microbiome is saliva, which contains bacteria shed from the oral surfaces and has been shown to represent the bacterial communities of the tongue, oral mucosa, teeth, tonsils, and throat (10).

Although the respiratory microbiota has received the most attention due to the main clinical manifestations of COVID-19, it is also important to consider the oral microbiota, as oral dysbiosis has been found to correlate with COVID-19 symptom severity and increased local inflammation (Soffritti et al. 2021) (11). It has been observed that the angiotensin-converting enzyme 2 (ACE2), the cell entry receptor of SARS-CoV-2, is expressed and therefore found in a wide variety of body tissues and fluids, including saliva (12,13) tongue or mouth floor (Xu et al. 2020) (14). Moreover, the salivary glands have been proposed as a potential target of SARS-CoV-2 infection, given the higher expression of ACE2 in the salivary glands compared to the lungs (15,16). However, there is limited knowledge about the interaction between SARS-CoV-2 and the oral microbiota, and whether the observed correlations are cause or consequence of the disease.

The objective of this study was to characterize the oral microbiota of children and adults and evaluate its relationship with age, household cohabitation, SARS-CoV-2 infection and COVID-19 severity during the pandemic lockdown in Barcelona, Spain.

## Methods

### Study design and population

The details of the study subjects and the procedures for sample collection and processing have been described in previous publications from researchers at the Hospital Sant Joan de Déu (HSJD) in Barcelona, Spain (17–19). These studies involved children and adults from the health region of the Metropolitan Area of Barcelona who volunteered to participate during the COVID-19 national lockdown period between April 28 and June 3, 2020.

### Inclusion and exclusion criteria

This analysis draws from a study conducted to understand, at the beginning of the confinement period, indoor transmission of COVID-19 from adult index cases to children. Participants living in family households were selected based on the following criteria: (i) at least one adult index case of SARS-CoV-2 infection confirmed by reverse transcription polymerase chain reaction (RT-PCR) at least 15 days prior to recruitment, and (ii) at least one child under the age of 15 living in the same household and subject to strict home confinement.

Inclusion was limited to participants who provided a saliva sample with a DNA concentration greater than 1 ng/µL.

### Definitions

Participants were classified into four age groups: <2 years, 2-5 years, 5-15 years, and adults. Children were defined as either SARS-CoV-2 infection cases or non-infected controls based on the results of RT-PCR and IgG/IgM antibody tests performed at the time of the household visit. Adult index cases of SARS-CoV-2 infection were classified as having a history of either severe or non-severe COVID-19 based on the requirement for hospitalization.

### Sample collection and processing

Saliva samples were collected during the first hour of the morning, before breakfast and tooth brushing, or at least one hour after the last food intake or last tooth brushing. Collection was performed using a micropipette for infants or by spitting into sterile tubes for older children and adults. Samples were subsequently transported to the Pediatric Biobank for Research at HSJD and stored at 80ºC (19) until they were sent for 16S rRNA gene sequencing at the FISABIO Sequencing and Bioinformatics Service in Valencia, Spain.

### DNA extraction and 16S rRNA gene sequencing

DNA from saliva samples was extracted using the MagNA Pure Compact Nucleic Acid Isolation Kit I (Roche, ref: 03730964001), according to the manufacturer’s instructions. The purified DNA was then used to amplify the V3-V4 variable regions of the 16S rRNA gene by polymerase chain reaction (PCR), using the following conserved primers: forward 5’-CCTACGGGNGGCWGCAG-3’ and reverse 5’-GACTACHVGGGTATCTAATCC-3’.

The PCR cycling conditions were as follows: 3 minutes at 95ºC, followed by 35 cycles of 30 seconds at 95ºC, 30 seconds at 55ºC, 30 seconds at 72ºC, then 5 minutes at 72ºC. DNA quantification was conducted using the Qubit 1X dsDNA HS Assay Kit (Thermo Fisher Scientific Inc., ref: Q33231). Only saliva samples with a DNA concentration greater than 1 ng/µL were sent for sequencing. 16S rRNA gene sequencing was performed in five separate runs on a MiSeq System (Illumina), using the MiSeq Reagent Kit v3 (Illumina), which generated 2×300 bp paired-end reads.

### Bioinformatic and statistical analysis

All analyses were performed with QIIME 2 (version 2023.9) (20), unless otherwise specified. A total of 65,371,819 reads were submitted for sequence quality control with DADA2 (21), using the *q2-dada2* plugin. To remove low-quality nucleotides, forward reads were trimmed at positions 10 and 280, and reverse reads at 10 and 230. These parameters were applied to all reads, except for those obtained from one of the sequencing runs, in which forward and reverse reads were trimmed at positions 10 and 275 and 10 and 225, respectively. After filtering, a total of 22,517,408 reads remained and were subsequently denoised into 15,949 amplicon sequence variants (ASVs). ASVs present in sequencing control samples were filtered out from the saliva samples to prevent contamination bias. As a result, one sample did not retain any reads after sequence quality control, and two others were excluded as they did not meet the sampling depth threshold set at 9,736 reads.

Diversity analysis was performed using the *q2-diversity* plugin at the ASV level rarefying at the same depth (9,736 reads) for each analysis. Alpha diversity was measured by Observed features and the Chao1 index (22) for species richness and the Shannon index (23) for species diversity. Beta diversity was measured using both weighted and unweighted UniFrac distance metrics (24,25), Bray-Curtis dissimilarity (26), and the Jaccard index (27). Additionally, Principal Coordinate Analysis (PCoA) (28) was performed using the distance matrices of all beta diversity metrics. Graphical representation included the beta diversity metric that better explained the composition differences.

Taxonomic classification was assigned to each ASV using the *q2-feature-classifier* plugin with a Naive Bayes classifier trained on the V3-V4 regions of the 16S rRNA gene, using reference sequences and taxonomy from the Greengenes2 2022.10 database (29).

Differential abundance analysis was performed using the Analysis of Compositions of Microbiomes with Bias Correction (ANCOM-BC) (30) method from the ANCOMBC package in R (version 4.3.2) (31), within the RStudio environment (version 2023.12.0 build 369) (32). Analysis was performed at both the ASV and genus levels. Features present in less than 10% of the total samples and taxa with a mean relative abundance of less than 0.01% were excluded.

Alpha and beta diversity metrics were compared across groups using the Kruskal-Wallis test (33) and Permutational Multivariate Analysis of Variance (PERMANOVA) (34), respectively, with Benjamini-Hochberg correction, using the *q2-diversity* plugin. Differences in taxonomic abundance between groups were assessed using the ANCOM-BC method with Holm-Bonferroni correction (35), via the ANCOMBC package in R. The adonis function from the vegan package (36) in R was used to estimate the proportion of variance explained by different metadata covariates. The total percentage of ASVs shared within households and between households was compared using the Wilcoxon rank-sum test (37), followed by a post hoc Dunn’s test (38). Statistical significance was defined as *p*-value < 0.05.

All plots and figures were generated in R using the following packages: *qiime2R* (39), *phyloseq* (40), *tidyverse* (41), *ggvendiagram* (42), and *pheatmap* (43).

## Results

### Characteristics of the study population

A total of 1465 individuals (381 adult index cases, 672 exposed children, and 412 exposed adults) from 381 households were initially considered for the present study (15). After application of the specific inclusion criteria, 142 adults and 265 children were included in the study, coming from 121 households with more than one participating family member (88 adults and 207 children). All 265 children were asymptomatic; however, SARS-CoV-2 infection was detected in 58 (21.89%) of them. All 142 adults had a prior confirmed SARS-CoV-2 infection, and 20 (14.08%) had undergone severe COVID-19 (Table 1).

**Table 1.** SARS-CoV-2 testing and COVID-19 hospitalization in the study population.

| Study group |  | Age |  |  | SARS-CoV-2 test result |  | Hospitalization due to COVID-19 |  |
| --- | --- | --- | --- | --- | --- | --- | --- | --- |
|  |  | Mean | SD | Range | Positive | Negative | Yes | No |
| Children<br>(n = 265) | Group 0<br>(aged <2 years,<br>n = 26) | 1.16 | 0.55 | 0.08-1.90 | 6 (23,08%) | 20 (76,92%) | - | - |
|  | Group 1<br>(aged 2-5 years,<br>n = 89) | 3.59 | 0.79 | 1.99-4.94 | 19 (21,35%) | 70 (78,65%) | - | - |
|  | Group 2<br>(aged 5-15 years,<br>n = 150) | 8.90 | 2.72 | 4.95-14.64 | 33 (22%) | 117 (78%) | - | - |
| Adults (n = 142) | Group 3<br>(n = 142) | 40.31 | 5.90 | 25.44-54.16 | (34,51%) | (65,49%) | 20 (14,08%) | 122 (85,92%) |

### 1. SARS-CoV-2 infection did not affect oral microbiota composition in asymptomatic, exposed children

The impact of SARS-CoV-2 infection on the oral microbiota was analyzed across different age groups of asymptomatic, exposed children. Among children under 2 years of age, no significant differences in bacterial richness or diversity were observed between SARS-CoV-2-infected and non-infected individuals (Figure S1A). Similarly, overall microbial community composition did not differ significantly between the two groups (Figure S1B). However, several taxa exhibited differential abundance at both the ASV and genus levels. Specifically, the genera *Granulicatella, Porphyromonas_A_859423, Bergeyella_A_791830* and *Haemophilus_D_736121* were enriched in SARS-CoV-2-infected children (Figure S2A). At the ASV level, two sequences classified as *Granulicatella ellegans*, two as *Prevotella melaninogenica*, and two as an unassigned *Haemophilus_D_736121* were enriched in SARS-CoV-2-infected children, whereas one sequence classified as *Rothia sp001808955* was enriched in non-infected children (Figure S2B).

Among children aged 2-5 and 5-15 years, no significant differences in bacterial richness or diversity were found between SARS-CoV-2-infected and non-infected individuals (Figure S3). However, within the 2-5 years age group, an ASV classified as *Haemophilus_A_sputorum* was enriched in non-infected children.

### 2. Severity of the COVID-19 disease’s symptoms alter the oral microbiota in adults

Saliva samples from the 20 adult index cases of SARS-CoV-2 infection with severe COVID-19 that required hospitalization exhibited lower bacterial richness and diversity compared to those from individuals with non-severe COVID-19 who did not require hospitalization (Figure 1A). Additionally, both groups differed significantly in terms of bacterial composition (Figure 1B), and several taxa showed differential associations at both the ASV and genus levels. Specifically, the genera HOT-345 (Human Oral Taxon 345), *Treponema_D*, F0422 (*Veillonella sp*.) and *Eubacterium_M* were enriched in individuals who did not require hospitalization, whereas an unassigned Lactobacillales, *Granulicatella, Veillonella_A, Streptococcus* and *Actinomyces* were enriched in individuals who required hospitalization (Figure 1C). At ASV level, two sequences classified as HOT-345 sp003260355 (ASV303 and ASV181 - *Clostridia*), one as *Filifactor villosus* (ASV437) and one as an unassigned *Nanoperiomorbus* (ASV815) were enriched in non-hospitalized individuals. In contrast, one sequence classified as an unassigned *Granulicatella* (ASV8), two as *Gemella sanguinis* (ASV43 and ASV40), one as *Oribacterium sinus* (ASV74) and one as *Streptococcus vestibularis* (ASV23) were enriched in hospitalized individuals (Figure 1D).

**Figure 1.**
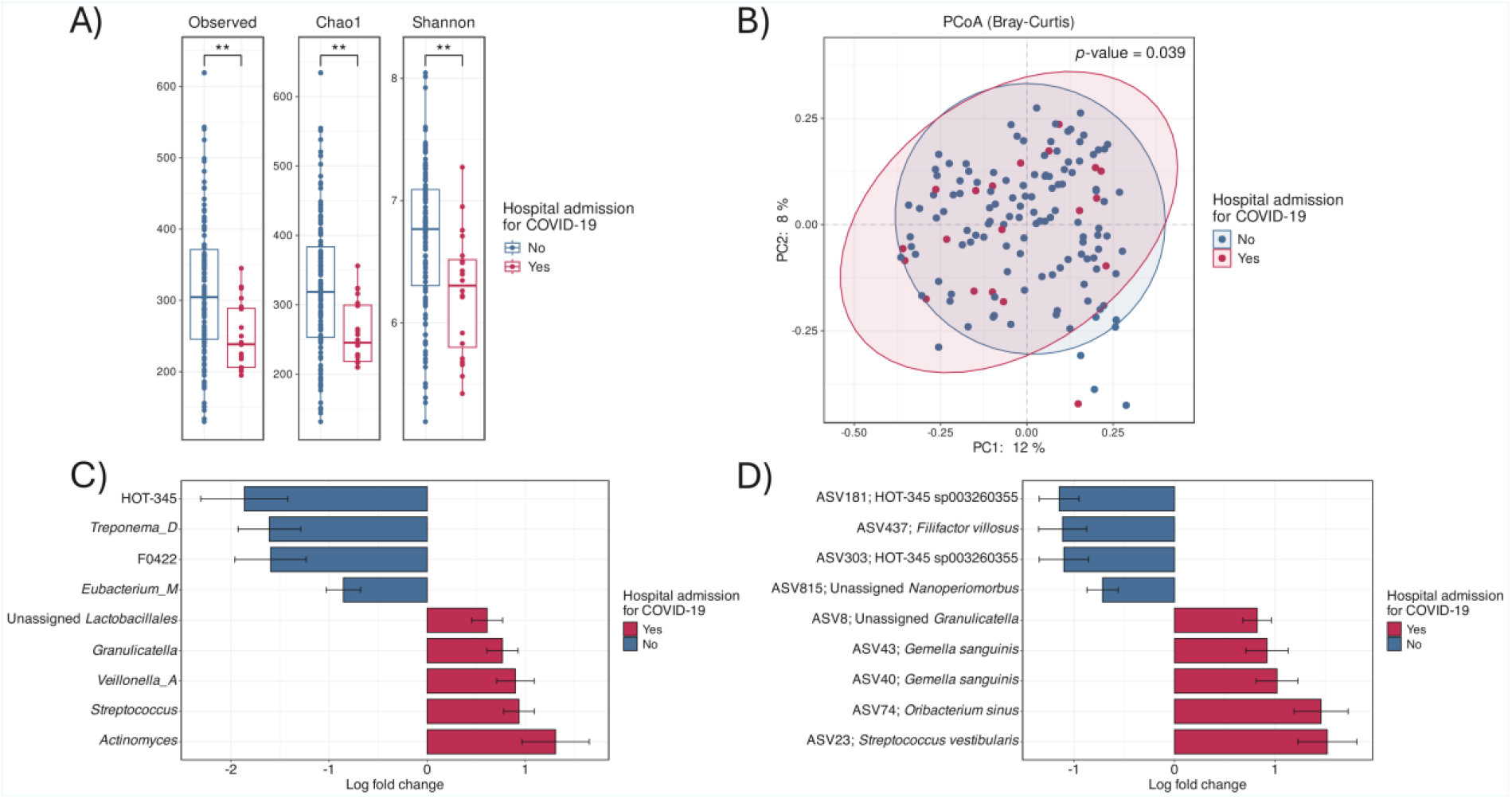
Diversity and differential abundance analyses of saliva samples from adults with confirmed SARS-CoV-2 infection who required hospitalization for COVID-19 (red) and those who did not (blue). Alpha diversity metrics are Observed features and the Chao1 index for bacterial richness and the Shannon index for bacterial diversity (panel A). *P*-values were calculated using the Kruskal-Wallis test with Benjamini-Hochberg correction. Significance levels are indicated as follows: “^****^” for *p*-values between 0 and 0.0001, “^***^” for *p*-values between 0.0001 and 0.001, “^**^” for *p*-values between 0.001 and 0.01, “^*^” for *p*-values between 0.01 and 0.05 and “ns” for *p*-values >0.05. Beta diversity is represented as a Principal Coordinate Analysis (PCoA) plot based on the Bray-Curtis dissimilarity distance matrix (panel B). *P*-value was calculated using PERMANOVA with Benjamini-Hochberg correction. Differential abundance analysis was performed using the ANCOM-BC method with Holm-Bonferroni correction and is represented at the genus level (panel C) and ASV level (panel D).

### 3. Oral microbiota is influenced by age

Alpha diversity analysis showed significant differences in bacterial richness and diversity across children age groups, with an observed increase with age. However, this increase was no longer significant between children aged 5-15 years and adults (Figure 2A).

**Figure 2.**
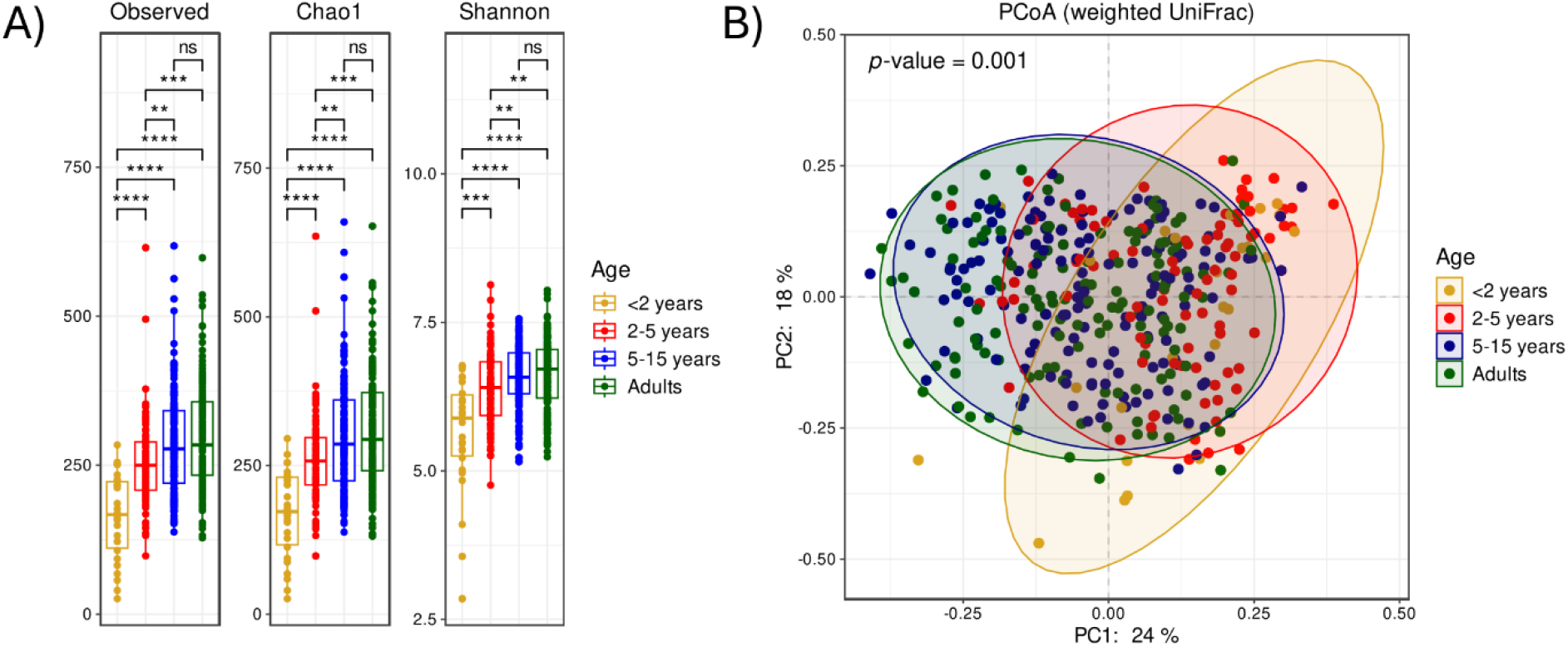
Diversity analysis of saliva samples across age groups. Alpha diversity metrics are Observed features and the Chao1 index for bacterial richness and the Shannon index for bacterial diversity (panel A). *P*-values were calculated using the Kruskal-Wallis test. Significance levels are indicated as follows: “^****^” for *p*-values between 0 and 0.0001, “^***^” for *p*-values between 0.0001 and 0.001, “^**^” for *p*-values between 0.001 and 0.01, “^*^” for *p*-values between 0.01 and 0.05 and “ns” for *p*-values >0.05. Beta diversity is represented as a Principal Coordinate Analysis (PCoA) plot based on the weighted UniFrac distance matrix (panel B). *P*-value was calculated using a PERMANOVA test with Benjamini-Hochberg correction.

**Figure 3.**
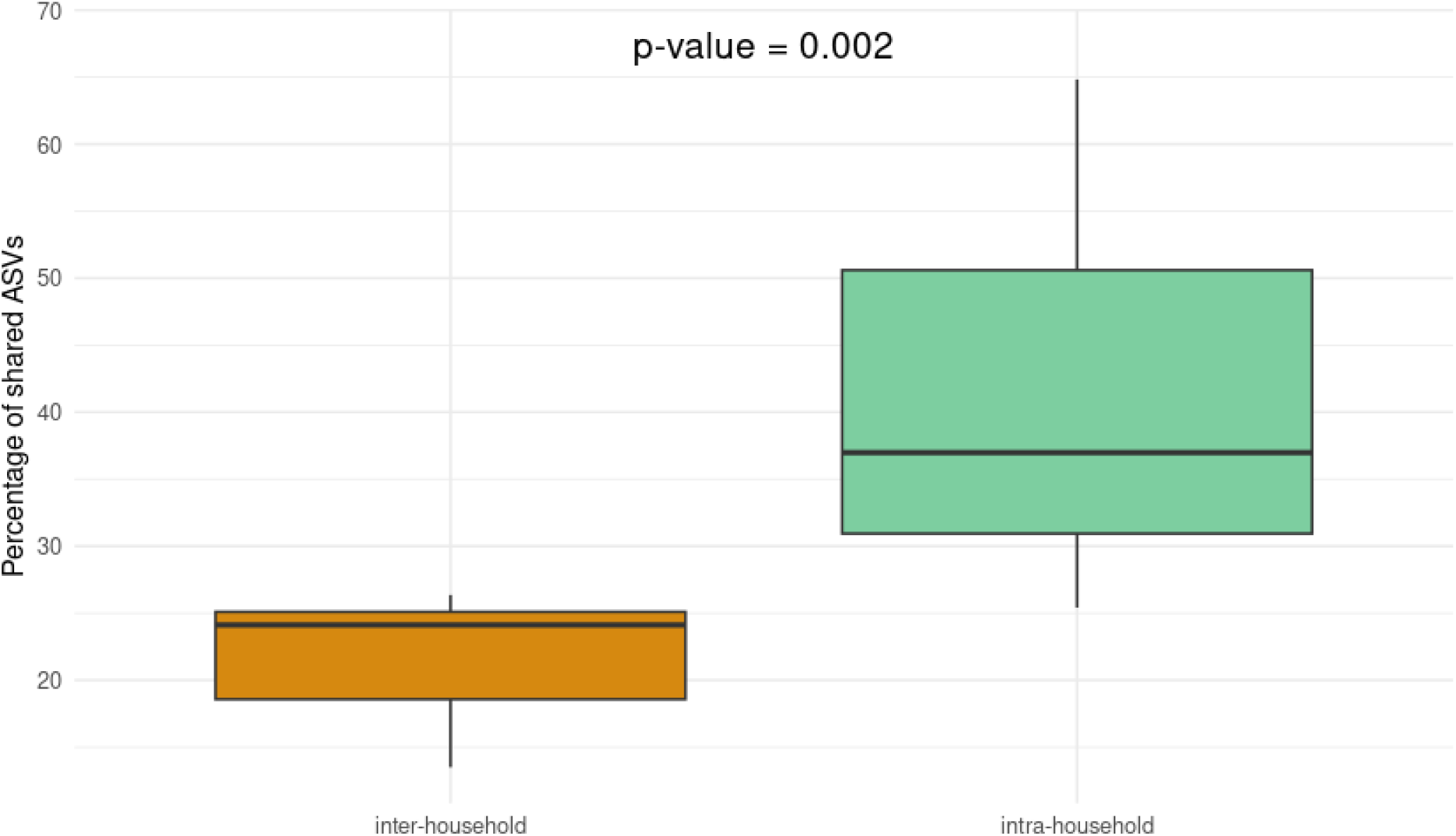
ASVs shared inter-families and intra-families. Boxplots represent the Percentage of shared ASVs (Y axis) between households (orange) and within household (green). Statistical significance among groups were inferred through Wilcoxon ranked sum test and post hoc Dunn’s test.

Beta diversity analysis also revealed differences in bacterial composition across age groups. Although all qualitative and quantitative metrics were statistically significant (PERMANOVA with Benjamini-Hochberg correction, *p*-value = 0.001), PCoA based on the weighted UniFrac distance matrix showed the highest percentage of variance explained by the First Principal Coordinate (PC1) (Figure 2B). Despite having a similar bacterial richness and diversity, the bacterial composition of children aged 5-15 years and adults differed significantly (PERMANOVA with Benjamini-Hochberg correction, *p*-value = 0.003).

To gain further insight, changes in bacterial composition within each age group were analyzed at the genus level, resulting in the identification of 263 different genera (Figure S4). Among these, 106 were present in children aged <2 years, 164 in children aged 2-5 years, 192 in children aged 5-15 years, and 220 in adults. There was a positive correlation between age and the total number of genera for each age group, as well as the number of unique genera: children aged <2, 2-5, 5-15, and adults exhibited 5, 7, 25, and 47 unique genera, respectively. Additionally, 93 genera were shared among all groups, representing 35% of the total number.

Analysis of the relative abundance of the genera revealed that a significant proportion exhibited values greater than 1%. Focusing on the 20 most abundant genera for each age group, *Streptococcus* and Neisseria were consistently among the most abundant genera across all groups (Figure S5). Interestingly, *Prevotella* showed an increasing trend with age, whereas *Granulicatella* and *Gemella* showed a decline. Supplementary Table 1 shows the relative abundance percentages for each genus across the different age groups.

### 4. Effects of household co-habitation on oral microbiota

Household co-habitation factor accounted for 69% of the variability in oral microbiota composition across all samples, as estimated using the Adonis function on the Weighted UniFrac distance matrix. Among 121 households with more than one participating family member, the percentage of shared ASVs within the same household (intra-household) ranged from 25.41% to 65.23% (median = 36.98%), which was significantly higher than the 13.44% - 26.36% range (median = 24.12%) observed between individuals from different households (inter-household) (Wilcoxon rank-sum test with Dunn’s correction, *p*-value = 0.002).

## Discussion

This analysis, drawing from a larger epidemiological study conducted in the midst of the initial lockdown period of the COVID-19 pandemic in Spain, was designed to identify changes in the oral microbiota according to SARS-CoV-2 infection, age and shared household. The alpha diversity metrics of the oral microbiota were observed to increase with age, which seemed to reach a plateau at some point between children aged 5-15 years (mean age 8.93±2.73) and adults (mean age 40.26±5.91), as no significant differences were observed between the two groups in any of the calculated metrics (Observed features, Chao1 and Shannon indices). These findings are consistent with those of other studies which have reported an increase in the alpha diversity of the oral microbiota with age during the first years of life (44,45,46). Particularly, Kahharova et al. (2020) characterized the salivary and dental plaque microbiomes of children at 3 time points: approximately 1 year old, 2.5 years old and 4 years old. They also collected saliva samples from the respective caregivers as well. Their results revealed an increment in alpha diversity metrics of both the salivary and dental plaque microbiomes with age. Significant changes in microbial composition were observed between 1 and 2.5 years of age, with less pronounced shifts up to 4 years old. However, both microbiomes did not reach the complexity observed in those of their caregivers. Holgerson et al. (45) conducted a longitudinal study with a similar objective to characterize the salivary microbiome in children from 2 days to 5 years of age. The results showed an increase in the alpha diversity of the salivary microbiome with age, with the most marked changes occurring between 3 and 18 months of age when the microbiome was still unstable.

This study also found a pronounced intra-household similarity in the oral microbiota composition (median = 36.98%) contrasting with the significant divergence detected between households (median = 24.12%). These results reinforce the emerging importance of the environment and social conditions in the shaping of the microbiota composition (47). They are also aligned with those reported by Valles-Colomer et al. (48) before the pandemic, showing that cohabiting individuals exhibited a median oral strain-sharing of 32%, whereas non-cohabiting individuals in the same population shared a median of 3%. The high intra-household oral microbiota similarity observed in our study may be attributed to the fact that it was conducted during the COVID-19 lockdown, a period characterized by prolonged and intensified cohabitation, and minimal extra-domiciliary contacts. However, all the datasets analyzed in the Valles-Colomer et al. (2023) study were retrieved in the pre-pandemic period. Additionally, the great intra-household similarity in our results could be also explained by the geographical restriction of our samples to the Metropolitan Area of Barcelona, as the Valles-Colomer et al. (2023) study included households from different cities, potentially increasing microbiota diversity across locations. Another reason might be that our study is based on an amplicon of the 16S rRNA gene, whereas the Valles-Colomer et al. (2023) study is analyzing WGS metagenomic data, which is a more precise approach to perform an estimation of the strain sharing rate. Similarly, Kort et al. (2014) documented that partners have a more similar oral microbiota composition than unrelated individuals (49). Furthermore, a higher degree of similarity between partners was associated with a higher frequency of kissing and a shorter period of time since the last kiss (49). Similarly, Song et al. (2013) found that cohabitation had a strong effect on oral, fecal, and especially skin microbiota composition (50). Moreover, the strongest effect was also observed between the parents, suggesting that cohabitation cannot overcome the differences due to age. Nevertheless, all of these studies have not been conducted in the context of strict home confinement, and therefore our data reflect an extreme situation with minimal contribution from exogenous microbiota.

As all adults in the study were confirmed SARS-CoV-2 cases by RT-PCR, this analysis focused on the impact of COVID-19 severity on the oral microbiota, showing that hospitalized patients exhibited significantly lower microbial richness and evenness, along with notable differences in bacterial composition compared to those who were not hospitalized. Although these results are consistent with previous reports of an inverse correlation between oral microbiota diversity and COVID-19 severity (11,51), some aspects, particularly the differential abundance findings, are challenging to interpret due to low relative abundances and limited data on certain genera such as HOT-345 and F0422, due to its recent discovery and taxonomic re-assignation. Nevertheless, some disease-associated bacteria appear to be congruent with previous work. Such is the case of *Granulicatella*, that we found associated at the genus and species level with COVID-19 severity, and that was reported by Wu et al (2021) (52) to be elevated in the oral cavity and the gut in individuals infected with SARS-CoV-2; or *Veillonella*, a biomarker of lactic acid levels, that was found associated with severe disease in our study and elevated in infected patients in the study by Soffritti et al. (2021). Although it has been reported that periodontal bacteria, which promote inflammation, are elevated in COVID patients compared to controls (50), we did not find them associated to disease severity. In relation to this, it is known that microbiota composition can be altered during and after infection by respiratory viruses (Grier et al. 2021) (53), and therefore differences in the length of time since infection clearance may represent an unaccounted confounded factor.

The main strength of this study lies in its implementation under strict home quarantine conditions, which ensured that all family members, regardless of age, had similar exposure to external factors influencing the composition of the oral microbiota. This approach minimized potential bias derived from differences in social interactions outside the home. Some limitations should also be taken into account for the interpretation of results. First, a convenience sample of participants was set up for the study on the basis of availability of samples. While convenience samples inherently carry a risk of selection bias, a clear inclusion criteria was defined to prevent arbitrary participant selection. Second, the study was conducted in a population characterized by specific sociodemographic and household conditions in a delimited geographic area during the early pandemic period. Any extrapolation of our findings to other geographic contexts or time periods should be made with caution. However, as discussed previously, our results of intra-household oral microbiota similarities were in agreement with those of other population studies conducted in diverse cities before the pandemic. Third, results regarding the adults’s hospitalization due to COVID-19, Moreover, it remains unclear whether the observed alterations in the oral microbiota were directly caused by the disease itself, and drugs received, particularly antibiotics, during hospitalization, given that the samples were collected when all the participants had recovered at home.

## Conclusions

During the COVID-19 pandemic, sharing a family household emerged as one of the primary determinants shaping the composition of the oral microbiota, underscoring the strong influence of close cohabitation on microbial community structure. Hospitalization, and to certain extent also the severity of the disease in SARS-CoV-2 infected adults, were associated with lower richness, diversity, and different oral microbiota composition compared to infected adults who did not require hospitalization. Asymptomatic SARS-CoV-2 infection did not have an important effect on the oral microbiota of children co-living with confirmed adults at any age range, suggesting a lack of direct antagonism between SARS-CoV2 and the oral microbiota.

## Data availavility

Datasets analysed in this study are publicly avalilabe in the public repository of European Nucleotide Archive at the European Bioinformatics Institute (EMBL-EBI), available at https://www.ebi.ac.uk/ena/browser/home under the study accession number PRJEB94431.

## Ethics approval

The study complies with the principles of the Declaration of Helsinki and the legal structure with respect to international human rights and biomedicine and the protection of personal data laws. The study was approved by the Ethics Committee of Fundació Sant Joan de Déu (PIC-59-20).

## Acknowledgements

The authors acknowledge the contributions of the Kids Corona study group members. In addition, the authors acknowledge to the Biobanc de l’Hospital Infantil Sant Joan de Déu per a la Investigació, which is integrated into the Spanish Biobank Network of Instituto de Salud Carlos III.

## Author’s contributions

Conceptualization, CM-A and PB; methodology, MB-F., GG-C, PB, AM, JJG-G, CL, RV, QB and MC; software, MB-F, GG-C, ALl, DH; Managed participant recruitment MFdS, RV, CL, MC; formal analysis, MB-F and GG-C; resources, CM-A, PB, QB and JJG-G; data curation, GG-C and MB-F; writing - original draft preparation, MB-F, GG-C and CM-A. Writing – review editing, PB, DH, MC, ALl, AM, QB, and RV. All authors have read and agreed to the published version of the manuscript. GG-C and MB-F contributed equally to this work.

